# gmos: Rapid detection of genome mosaicism over short evolutionary distances

**DOI:** 10.1101/053694

**Authors:** Mirjana Domazet-Lošo, Tomislav Domazet-Lošo

## Abstract

Prokaryotic and viral genomes are often altered by recombination and horizontal gene transfer. The existing methods for detecting recombination are primarily aimed at viral genomes or sets of loci, since the expensive computation of underlying statistical models often hinders the comparison of complete prokaryotic genomes. As an alternative, alignment-free solutions are more efficient, but cannot map (align) a query to subject genomes. To address this problem, we have developed gmos (Genome MOsaic Structure), a new program that determines the mosaic structure of query genomes when compared to a set of closely related subject genomes. The program first computes local alignments between query and subject genomes and then reconstructs the query mosaic structure by choosing the best local alignment for each query region. To accomplish the analysis quickly, the program mostly relies on pairwise alignments and constructs multiple sequence alignments over short overlapping subject regions only when necessary. This fine-tuned implementation achieves an efficiency comparable to an alignment-free tool. The program performs well for simulated and real data sets of closely related genomes and can be used for fast recombination detection; for instance, when a new prokaryotic pathogen is discovered. As an example, gmos was used to detect genome mosaicism in a pathogenic *Enterococcus faecium* strain compared to seven closely related genomes. The analysis took less than two minutes on a single 2.1 GHz processor. The output is available in fasta format and can be visualized using an accessory program, gmosDraw (freely available with gmos).

## Introduction

Similar to viruses, prokaryotes often exchange homologous genetic material by horizontal gene transfer (HGT). Such sequences newly acquired from other organisms may help the host to adapt to environmental changes [1–3], or become resistant to drugs [1,2]. Most existing methods for the detection of genome mosaicism have been developed for the analysis of viral genomes or sets of prokaryotic genes. For instance, there is a range of programs designed for the analysis of recombination in HIV-1 [4,5], HCV [6,7], and HBV [8]. There is also a group of programs based on statistical or probabilistic methods for modeling recombination events between bacterial genomes, e.g., RDP4 [9], BratNextGen [10], ClonalFrame [11], ClonalFrameML [12], ClonalOrigin [13], fineSTRUCTURE [14], and orderedPainting [15]. However, these methods typically handle only a subset of genomic loci, since the analysis of complete genomes is hindered by computationally expensive modeling processes and the requirement of multiple sequence alignment as input. Although efficient general purpose tools for local pairwise alignments that handle complete genomes are available, e.g., BLASTN [18] and Mega BLAST [19], they are not directly applicable for detecting recombination, as they tend to maximize extension of the local alignment along every subject genome without considering alignment information from other subject sequences.

As an efficient alternative, alignment-free programs can be used for annotating genome mosaic structure in whole bacterial genomes. An example is alfy, which was applied to *Escherichia coli* and *Staphylococcus aureus* genomes affected by horizontal gene transfer [16,17]. However, an alignment-free program cannot map query to subject regions, i.e. it cannot provide information about the exact positions of matches between a query and a subject.

To address this problem of efficient and informative detection of genome mosaicism in closely related prokaryotes, we have developed a new program, gmos. This program analyzes one or more query genomes (either complete or draft) when compared to a set of *n* subject genomes. The query mosaic structure is determined directly from the efficient pairwise and, only when necessary, multiple sequence alignment of the query and subject genomes.

We show that gmos is a very efficient tool for detecting genome mosaic structure; its speed and memory performance are comparable to alignment-free solutions. We also demonstrate its accuracy on simulated sequences by comparing it to a statistically based tool that models recombination events, and by analyzing the recombinant structure of the pathogen *Enterococcus faecium*. gmos was written in C programming language and is available under the terms of the GNU General Public License from http://www.zpr.fer.hr/osobe/mirjana/gmos/. The software package includes documentation and test data.

## Implementation

The input to gmos are two files: the first file comprising *m* query genomes and the second file comprising *n* subject genomes (in both files, input sequences can be either complete or draft). The program compares each query sequence to all subject sequences and outputs its mosaic structure. For the sake of simplicity, the implementation details are further presented only for the case when a single query genome is compared to a set of *n* subject genomes.

## Overview of gmos

Let *Q* be a query DNA sequence and ***S*** be a set of DNA subject sequences: ***S*** = {*S*_*1*_, *S*_*2*_, ¨, *S*_*n*_} represented as strings over the alphabet Σ = {A, C, G, T}. gmos takes *Q* and ***S*** as input and computes the mosaic structure of *Q*, denoted *M. M* represents a list of best local alignments between regions of *Q* and regions from any subject of ***S***. In the internal representation of *Q* and ***S***, the characters that are not elements of the alphabet Σ are masked and treated as mismatches in the alignment computation.

There are two main phases of the program (Fig 1): (i) the construction of the set ***L***, which comprises *n* lists of local alignments between *Q* and each *S*_*i*_, *i* = 1, ¨, *n*, and (ii) computation of the mosaic structure of the query, *M*. The program returns *M*, a list of local alignments between query regions and best locally aligned subject regions, each at least *f* characters long (default: 200 bp).

**Fig 1.**
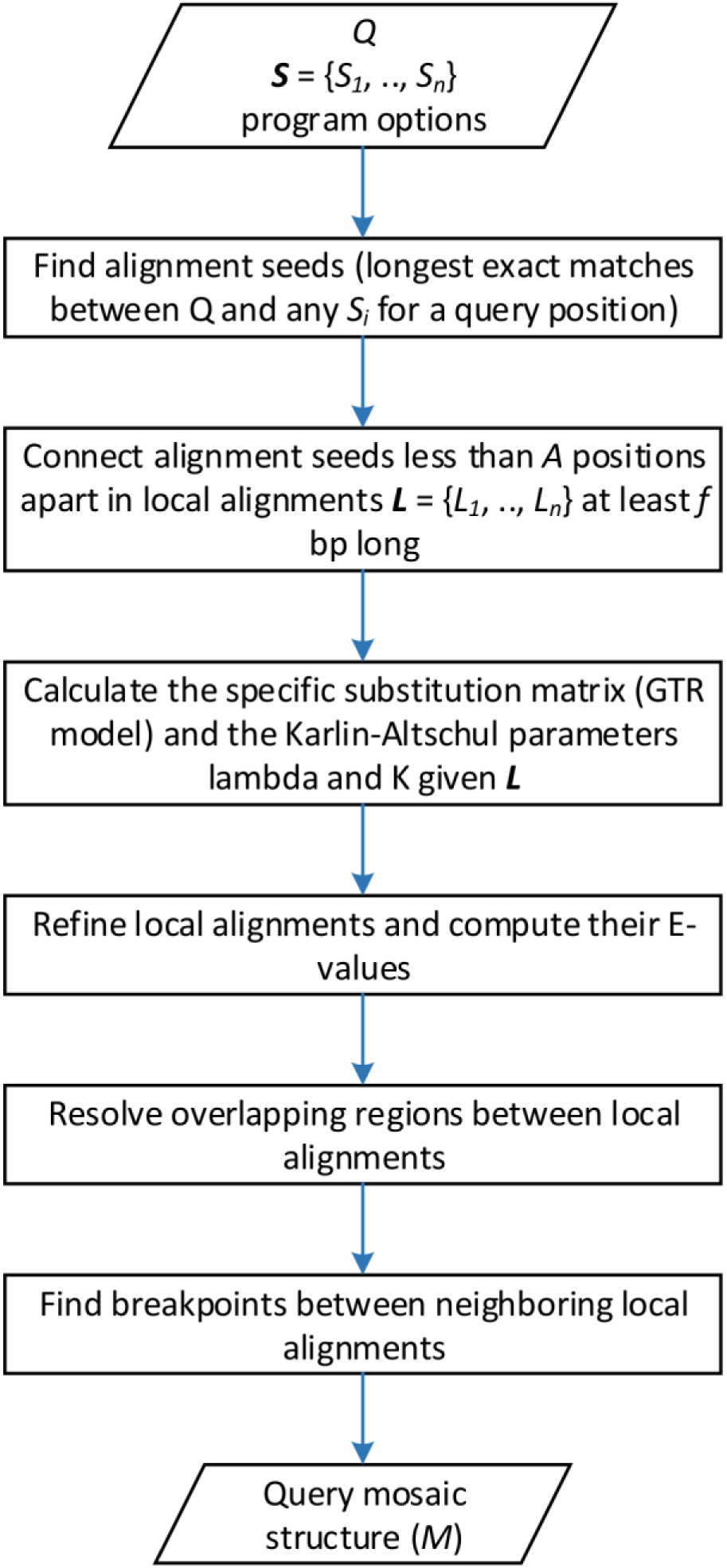
Overview of gmos workflow. The program computes a mosaic structure (*M*) of a query sequence (*Q*), when compared to a set of *n* subject sequences ***S*** = {*S*_*1*_, *S*_*2*_, ¨, *S*_*n*_}. ***L*** is the set of lists *L*_*i*_, where each *L*_*i*_ is the list of local alignments between *Q* and *S*_*i*_. To compute the query mosaic structure (*M*) local alignments from ***L*** are processed to determine the most similar subject regions for each query region.

In the first phase, local alignments between a query (*Q*) and each subject *S*_*i*_, *i* = 1, ¨, *n*, are constructed using their exact matches as alignment seeds. The minimal length of an exact match, considered to be an alignment seed, is computed for a pair (*Q*, *S*_*i*_) from the length of the shortest unique substring (or shortly, shustring [27]) for each query position. The exact matches (shustrings) between the pairs (*Q*, *S*_*i*_), *i* = 1, ¨, *n*, are determined using an enhanced suffix array [22–24] constructed of all query and subject genomes. An exact match between *Q* and *S*_*i*_, longer than expected by chance alone [25] and longer than any other exact match between *Q* and *S*_*j*_, *j* ≠ *i*, which starts at the same query position, is stored as an interval in the corresponding interval tree of a (*Q*, *S*_*i*_) pair. The interval trees are constructed for each pair (*Q*, *S*_*i*_), similarly to the procedure used in [16]. The regions between the close-enough intervals in an interval tree are aligned using the Needleman-Wunsch algorithm [26] and then connected in a single local alignment. Close-enough intervals are exact matches at most *A* bp apart (*A* is set to 70 bp by default, but can be adjusted by the user). The first phase results in the set of *n* lists of local alignments (the set ***L***).

In the second phase, the mosaic structure of a query genome, *M*, is constructed from *n* lists of local alignments (the set ***L***). For each query region locally aligned to more than one subject region, the most similar subject region is chosen by comparing the pairwise alignment scores of each query-subject pair. In addition, efficient multiple sequence alignment is applied to detect the best local alignment(s) when multiple subjects are locally aligned to overlapping query regions. We apply a version of multiple sequence alignment called the central star approach [31, 32], where the query is a central sequence to which the multiple subjects are locally aligned and thus locally aligned between themselves. The procedure is very fast, since we construct the multiple sequence alignment only over short overlapping regions and the subject regions were already pairwise aligned to query in the first phase. *M* finally comprises a list of local alignments at least *f*bp long (default: 200), where each local alignment is the best alignment between a query region and the overlapping subject regions. Also, the *E*-value of each local alignment is computed as in BLAST [27,28]: *E* = *Kmn* · e^−λ*S*^ where *λ* (lambda) and *K* are the Karlin-Altschul parameters [27,28], *m* is the query length, *n* is the total length of all subject sequences and *S* is the new score of a local alignment recomputed using the refined DNA substitution scoring matrix (GTR model) [29]. This DNA scoring matrix is computed from the initial alignments in ***L***. The parameters *λ* and *K* are analytically computed as previously described for ungapped alignment [27,28,30], given that in the case of closely related sequences, the values of *λ* and *K* for gapped alignments are expected to be similar to *λ* and *K* for ungapped alignments [31].

## Results and Discussion

### Runtime and memory analysis

The running time and memory analysis of gmos was performed on closely related sequences simulated using Dawg [32] and compared to an alignment-free tool, alfy [16]. We chose alfy for comparison because it uses a similar indexing approach to gmos for computing exact matches and therefore is the best reference for estimating the alignment construction efficiency of gmos. Each data set used in the analysis comprised a query and *n* subject sequences of equal length: the sequence lengths ranged from 10 kb to 5 Mb and *n* ∈ {10, 50, 100}. Both programs were run with their default settings and the running time for each graph point was calculated as the average over 10 runs (each on a different data set with the same characteristics). Figure 2 shows that gmos is similar to alfy regarding running time and memory consumption. For example, the gmos comparison of a query sequence to the data set of 100 subject sequences of total length 500 Mb took 17 minutes and used 11.4 GB of memory, while alfy took 21 minutes and required 11.3 GB of memory (Fig 2). This analysis also showed that the running time and memory consumption of gmos are approximately linear for the tested parameters. This and all subsequent analyses were performed on a single processor (Intel Xeon E5-2620v2, 2.1GHz).

**Fig 2.**
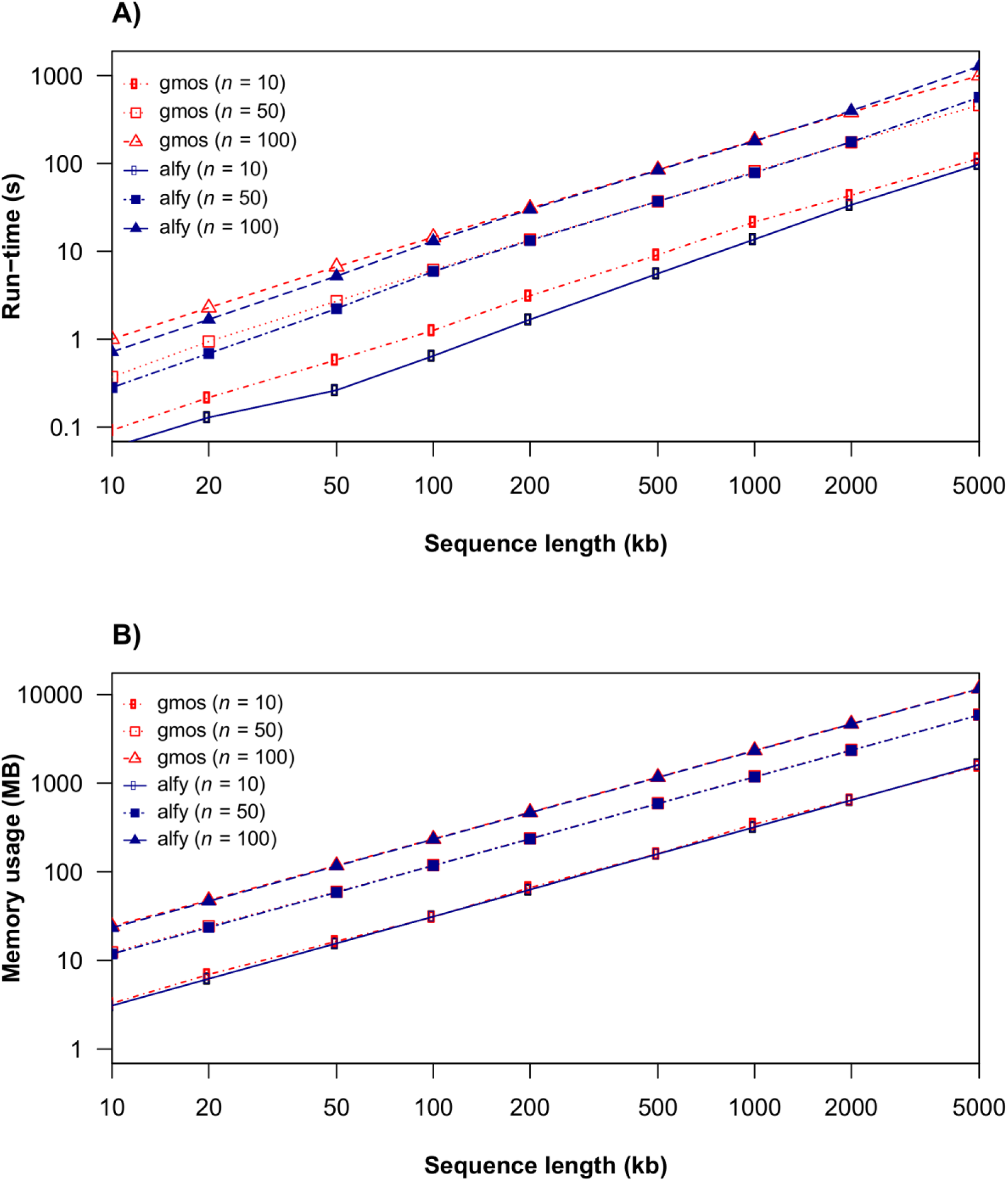
Run time and memory usage of gmos. A) Run times and (B) memory usage of gmos and alfy as a function of sequence length and the number of subject sequences, *n*.

### Accuracy of gmos

Next, we tested gmos for its accuracy on simulated data sets consisting of a query and three subject sequences (*S*_*1*_, *S*_*2*_, *S*_*3*_). All sequences were of equal length, *L*, where *L* ∈ {10 kb, 25 kb}. The data sets were simulated using Dawg [32] and the underlying phylogeny (Fig 3) was drawn using MEGA6 [33]. The query was a recombinant constructed as a concatenation of 5 equally long segments (the length of each segment was *L*/5). The closest subject sequence of each query segment switched between *S*_*1*_ (Fig 3A, genealogy A) and *S*_*2*_ (Fig 3B, genealogy B) in the following order: *S*_*1*_- *S*_*2*_- *S*_*1*_- *S*_*2*_- *S*_*1*_ (as previously described in [16]). The evolutionary distance (the relative number of segregating sites, *s*) between a query segment and the immediate common ancestor between a query and its most closely related subject sequence (either *S*_*1*_ or *S*_*2*_) ranged from 0.001 to 0.11 (Figs 3 and 4). The evolutionary distance from the query’s closest subject sequence (switching between *S*_*1*_ and *S*_*2*_) to the common ancestor was also *s*. Accuracy was measured as the fraction of query nucleotides correctly assigned to the true closest relative (either *S*_*1*_ or *S*_*2*_). Each graph point represents the average accuracy over 10 runs (Fig 4). We used a strict approach in which the following three cases were all considered as inaccurate results: (i) a false subject was returned; (ii) no subject was returned; (iii) a best hit was not singled out (i.e. both *S*_*1*_ or *S*_*2*_ were returned as best hits). The highest accuracy was obtained for *s* = 0.1 and *L* = 10 kb (0.94 ± 0.02) and for *s* = 0.05 and *L* = 25 kb (0.96 ± 0.03) (Fig 4). As expected, the program’s accuracy declines for *s* > 0.1, since it was calibrated for detecting recombination between closely related genomes or for detecting horizontal gene transfer where compared sequences are relatively similar.

**Fig 3.**
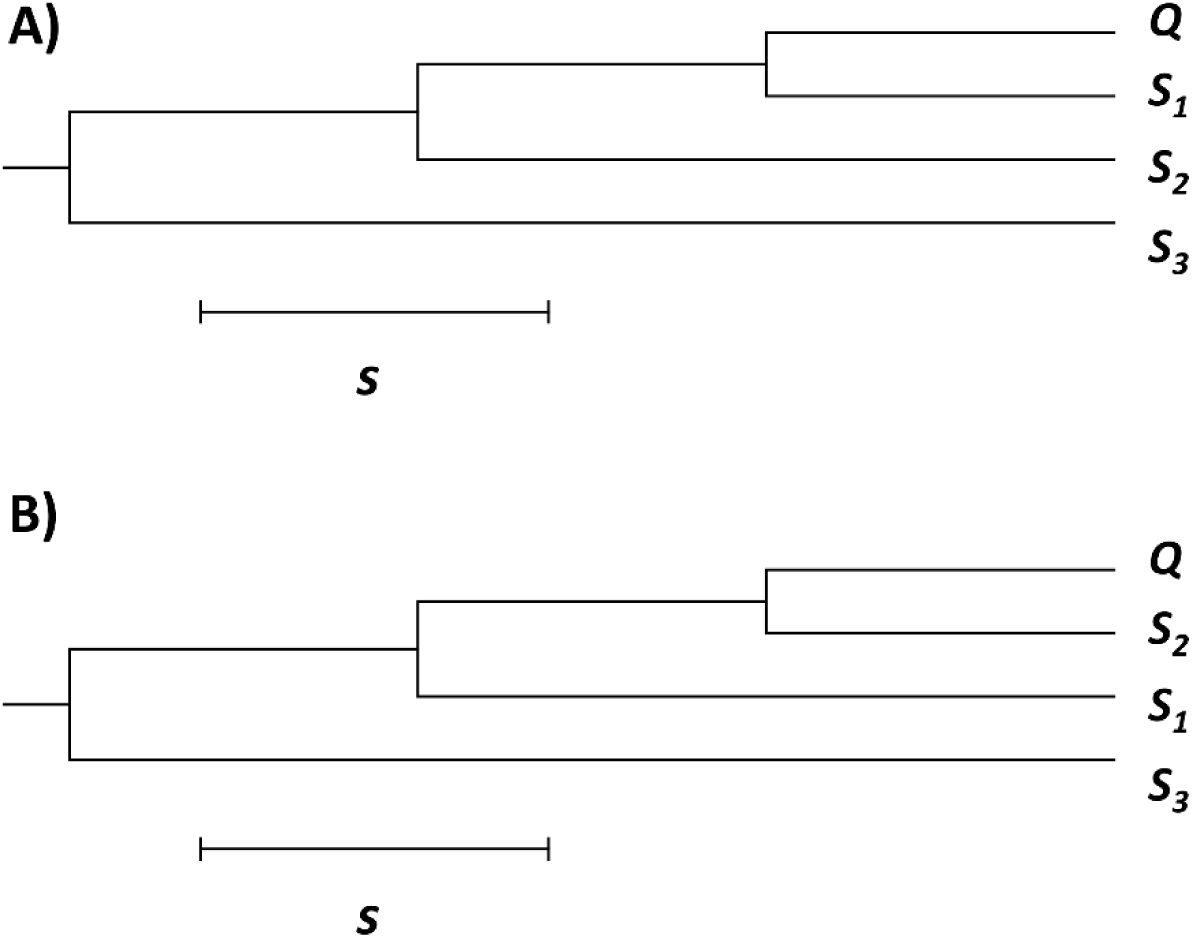
Genealogies of a query sequence *Q* and the subject set {*S*_*1*_, *S*_*2*_, *S*_*3*_}. *Q* is a recombinant comprising 5 segments, whose genealogy alternates between A and B in order: A - B - A - B - A. The evolutionary distance (the relative number of segregating sites) is denoted *s*. (A) Genealogy A: *Q* is most closely related to *S*_*1*_. (B) Genealogy B: *Q* is most closely related to *S*_*2*_.

**Fig 4.**
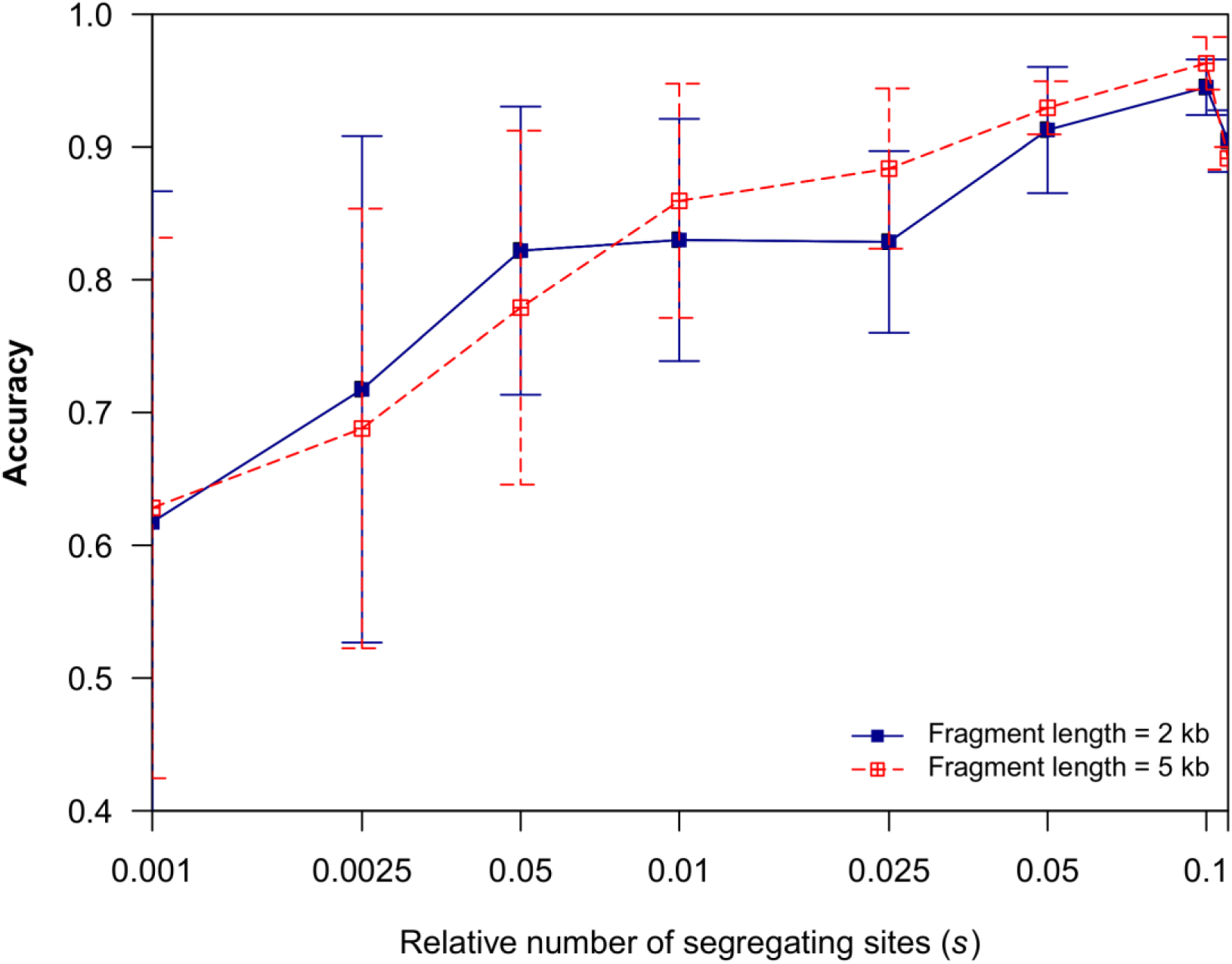
Accuracy analysis of gmos as a function of evolutionary distance. The relative number of segregating sites between a query segment and its immediate ancestor, *s*, is also the distance between the query’s immediate ancestor and the query’s closest relative among subject sequences (as illustrated in Fig 3).

In the next experiment using simulated data (generated using Dawg [32]), a query sequence (*Q*) was compared to a set of 10 subject sequences (*S*_*1*_, *S*_*2*_,…, *S*_*10*_), each 10 kb long. In the experiment, sequences *S*_*1*_ and *S*_*2*_ were identical and shared the immediate common ancestor with a query sequence (S1 Fig.). In this example, *S*_*1*_ and *S*_*2*_ were the closest relatives of *Q* along its entire genome. The analysis was conducted for evolutionary distance (the relative number of segregating sites), *s*, ranging from 0.001 to 0.1 (S2 Fig.). Accuracy was calculated by the fraction of query nucleotides correctly assigned to *S*_*1*_ and *S*_*2*_. For each evolutionary distance, the graph point represents the average accuracy over 10 runs (S2 Fig.). The analysis showed that both *S*_*1*_ and *S*_*2*_ sequences returned the same nucleotides along the query genome. The highest accuracy was achieved for *s* = 0.05 (0.98 ± 0.02). Again, since the program has been developed for closely related sequences, accuracy of the program declined for *s* > 0.1 (S2 Fig.), similar to the result of the previous experiment (Fig 4).

### Comparison to other tools

We also compared gmos to a statistically based software, ClonalOrigin [13], on simulated and real data (S1 and S2 Files). We chose ClonalOrigin for this comparison because it is a popular statistically based tool used for determining bacterial recombination [15] that returns recombining sequences (both donor and recipient sequences) and their positions within the alignment, therefore allowing comparison of similar parameters.

In this experiment, gmos was run with the default options. ClonalOrigin was run by following the program’s guidelines [13]. In case of simulated data, the running time (*t*), the rate of true positives (*TPR*), and the rate of false positives (*FPR*) of both programs were calculated as the average over 10 runs (each run on a different data set with the same characteristics) (S1 File). The simulated data sets were generated using Dawg [32], where each data set was comprised of four sequences: a recombinant query sequence and three subject sequences (*S*_*1*_, *S*_*2*_, and *S*_*3*_), each 10 kb long. A query was constructed to be most closely related to *S*_*1*_, with its second and fourth segment acquired from *S*_*2*_, as described in the section above (Fig 3), thus simulating the two recombination events. The data sets were simulated over a range of evolutionary distances (the relative number of segregating sites), *s*, from 0.001 to 0.1. The running time, the true positive rate (*TPR*), and the false positive rate (*FPR*) were measured for each *s* value (S1 File). *TPR* represents the fraction of the query positions correctly detected to be horizontally transferred from *S*_*2*_, and *FPR* represents the fraction of the query positions incorrectly detected to be horizontally transferred from *S*_*2*_. In case of gmos, *TPR* and *FPR* are the fractions of query positions assigned to *S*_*2*_, where *S*_*2*_ was recognized as the best hit, i.e. the most closely related subject sequence. In case of ClonalOrigin, *TPR* and *FPR* were calculated only on returned recombination events where the donor sequence was *S*_*2*_ and the query was the recipient.

Both programs had lower sensitivity over very small evolutionary distances, *s* = 0.001 to 0.0025 (S1 File). At these distances, subject sequences *S*_*1*_ and *S*_*2*_ were both very closely related to each other and to the query, so neither program could detect recombination well; gmos often returned both subjects *S*_*1*_ and *S*_*2*_ as the best hits for a particular query region, and ClonalOrigin could not determine one or both recombination events. The sensitivity of both programs improved for larger evolutionary distances (*s* ≥ 0.005), with ClonalOrigin surpassing gmos over moderate evolutionary distances (*s* = 0.005 to 0.025) and gmos outperforming ClonalOrigin over larger evolutionary distances (*s* ≥ 0.05).

In the case of gmos, *TPR* increased and *FPR* decreased with evolutionary distance and the program performed best for the highest evolutionary distance, *s* = 0.1 (*TPR* = 94.42%, *FPR* = 1.64%). In comparison, ClonalOrigin achieved the best result for *s* = 0.01 (*TPR* = 91.78%, *FPR* = 5.04%). However, in the case of ClonalOrigin, *FPR* and the number of incorrectly detected recombination events increased with larger evolutionary distances, *s* > 0.025. The program was not tested for *s* > 0.05 due to the substantial decreases in performance and computational burden at these values. In addition, gmos outperformed ClonalOrigin by orders of magnitude in terms of speed for all data sets; analysis of a single data set by gmos took around 0.04 seconds across all evolutionary distances, while the running time of ClonalOrigin increased with larger values of *s* (S1 File), ranging from 31 minutes for *s* = 0.001 up to 3.4 hours for *s* = 0.05.

Finally, gmos and ClonalOrigin were used for analysis of a recently sequenced pathogen strain, *Enterococcus faecium* 1,231,408 (2.8 Mb), which was compared to seven other *Enterococcus faecium* genomes (in total: 20 Mb; listed in Table 1). A previous study [20] revealed the strain’s recombinant nature; although classified as an *Enterococcus faecium* clade A strain, it comprises two major gene clusters acquired from clade B strains that span almost 25% of its genome. The output of both programs for this dataset (S2 File) were consistent with the previously reported results [20].

**Table 1.** Enterococcus faecium strains used in the analysis

| GenBank Accession Number | Strain | Short name | Clade |
| --- | --- | --- | --- |
| ACBB000000000 | 1,231,408 | 408 | **A** |
| ACBA000000000 | 1,231,410 | 410 | **A** |
| ACAY000000000 | 1,231,501 | 501 | **A** |
| ACAX000000000 | 1,231,502 | 502 | **A** |
| ACAS000000000 | 1,230,933 | 933 | **A** |
| ACAZ000000000 | 1,141,733 | 733 | **B** |
| ACBC000000000 | Com12 | Com12 | **B** |
| ACBD000000000 | Com15 | Com15 | **B** |

As a prerequisite for the ClonalOrigin analysis, each strain’s contig was aligned to contigs of seven other *Enterococcus faecium* genomes (Table 1) using Mugsy [34]. The multiple sequence alignment procedure resulted in 21 alignment blocks for the strain’s supercontig 1.3 and 13 alignment blocks for the strain’s supercontig 1.5. Each alignment block represented multiple sequence alignment of the eight strains. ClonalOrigin analyzed each alignment block separately. The results revealed horizontal gene transfer from clade B strains to genes of both supercontigs (S2 File), as previously reported [20]. The analysis of an alignment block lasted between 11 minutes and 35.9 hours, depending on the size of the block and the number of recombination events detected. However, these values are minimal estimates as the program did not converge in the last phase of the analysis where global parameters are applied.

In contrast, gmos was applied directly to whole genomes (no preparation of the data required). The recombinant strain (*Enterococcus faecium* 1,231,408) was treated as a query and the other seven genomes were considered subjects. In accordance with the original result [20], gmos detected horizontal gene transfer from strains of clade B to supercontigs 1.2, 1.3, and 1.5 of *Enterococcus faecium* 1,231,408; gmos analysis showed that the strain’s supercontig 1.3 is most similar to the supercontigs 1.2 of two clade B strains, *Enterococcus faecium* Com12 and 1,141,733 (Fig 5, S2 File), as previously shown. The gmos direct analysis of whole genomes took 70 seconds, around 500 times faster than ClonalOrigin, and required 600 MB of memory. Taken together, these results show that both gmos and ClonalOrigin are applicable in real situations with gmos being superior in terms of speed, input data size, and usage of raw sequences.

**Fig 5.**
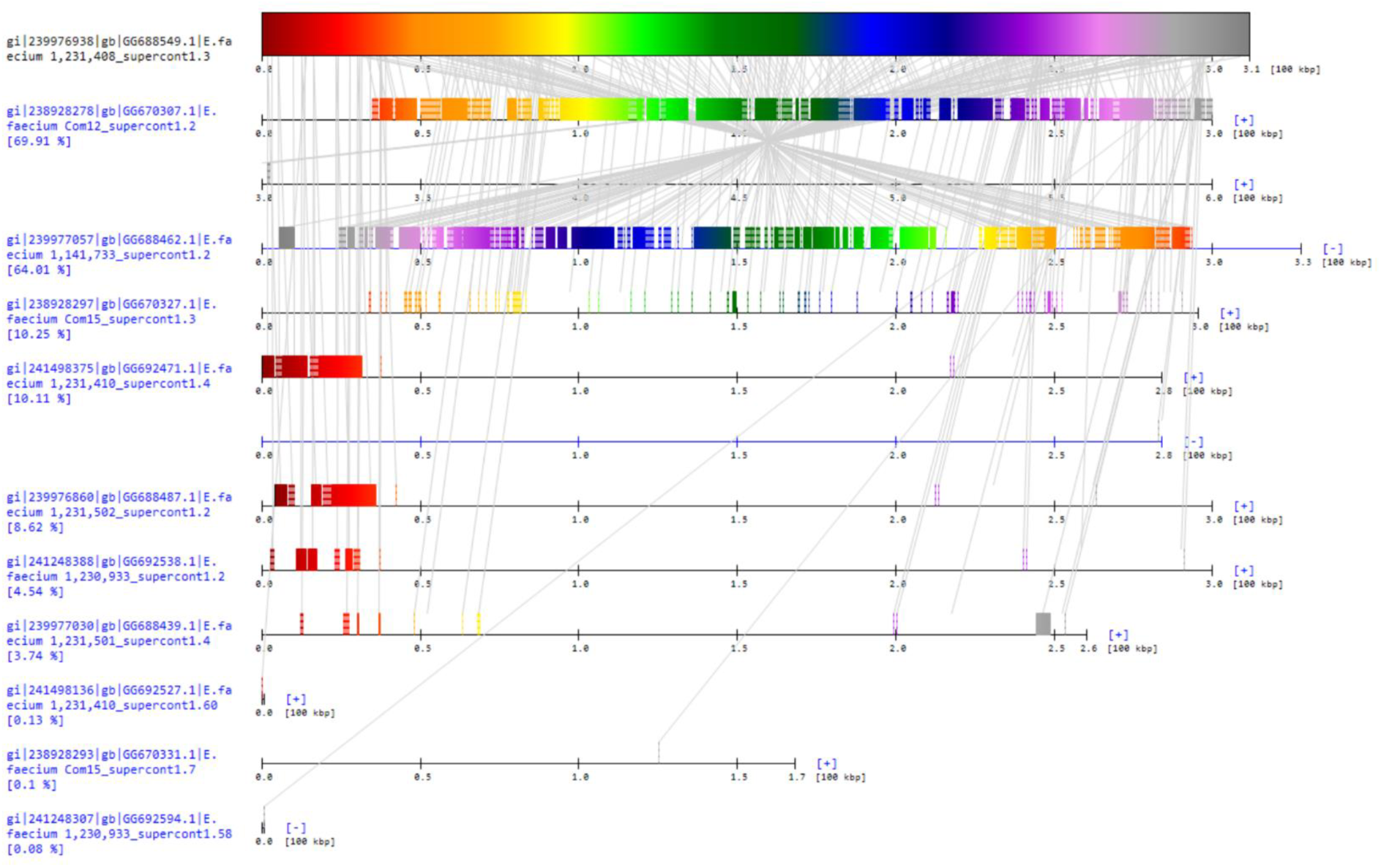
gmos analysis of *Enterococcus faecium* 1,231,408, supercontig 1.3. The results of gmos analysis of the strain *Enterococcus faecium* 1,231,408, supercontig 1.3, in comparison to the set of 7 subject *Enterococcus faecium* genomes comprised of multiple contigs. In accordance with the previous results, gmos analysis revealed the recombinant nature of the query strain: supercontig 1.3 is most similar to supercontigs 1.2 of the strains Com12 and 1,141,733, both of clade B, while the query strain itself is from clade A. The results are drawn using the accessory tool, gmosDraw.

## Availability

gmos was written in C programming language and runs under the Unix/Linux operating system. Its source code is available under the GNU General Public License. gmosDraw was written in C# programming language and runs under Windows operating system. Both programs are freely available at: http://www.zpr.fer.hr/osobe/mirjana/gmos/. The software packages include test data and documentation for installing and running the programs.

## Conclusion

gmos rapidly detects a mosaic structure of a query genome when compared to a set of whole genomes (either complete or draft), and does not require pretreatment of input sequences. These properties make it an ideal tool for situations where rapid screens for recombination are needed and as a complement to more precise, statistically based programs. Its output is the list of best locally aligned regions in multi-fasta format. Additionally, a sequence mosaic structure can be visualized using the accessory program, gmosDraw, which enables easy interpretation of the results.

## Acknowledgments

We thank Robert Bakarić, Marko Madunić, and Kristian Vlahoviček for helpful comments on the manuscript.

---

**Supporting Information**

**S1 Fig.**
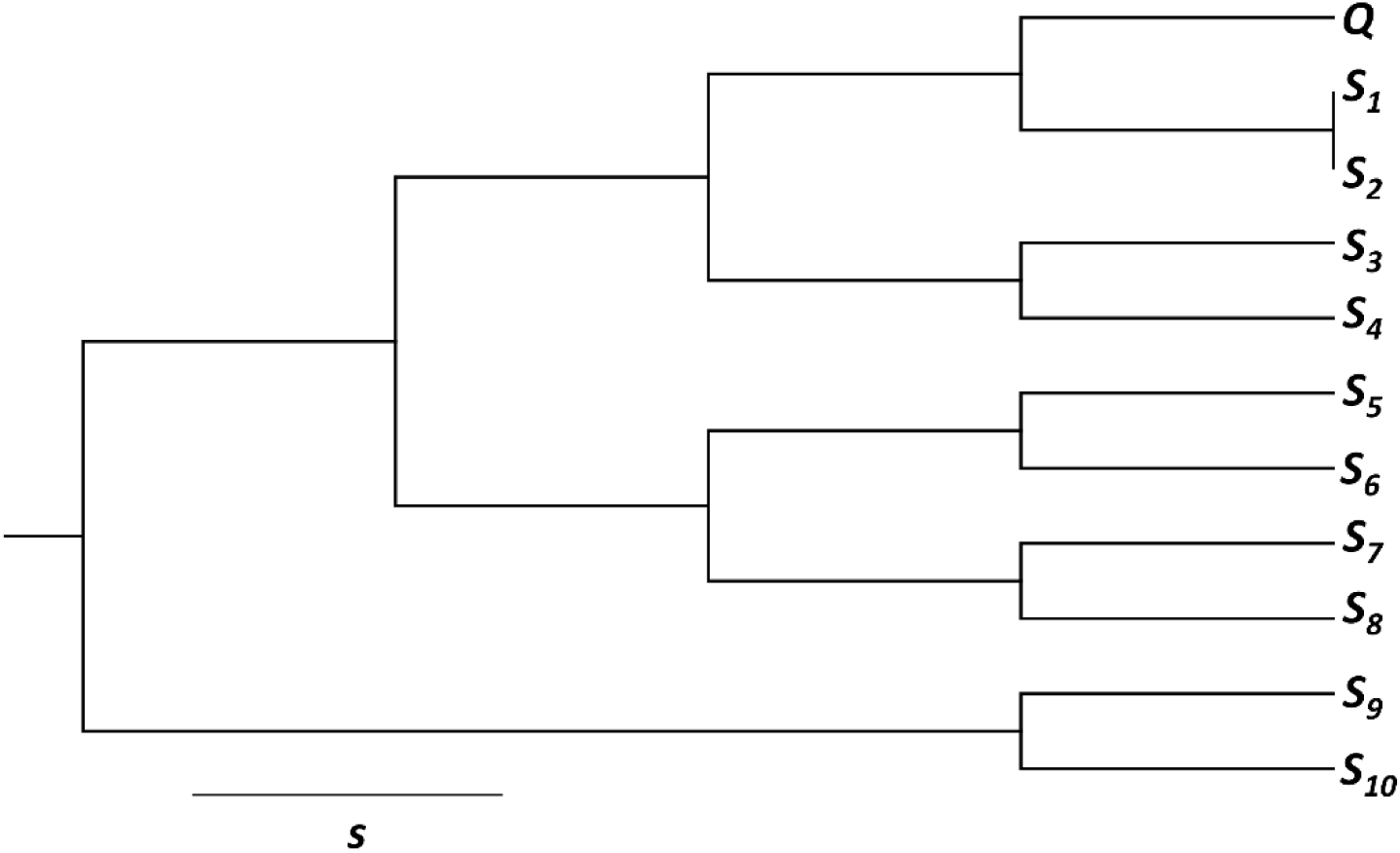
Genealogy of subject sequences where *S*_*1*_ and *S*_*2*_ are equally closely related to a query. Genealogy of a query sequence *Q* and the subject set {*S*_*1*_, *S*_*2*_,…, *S*_*10*_} (simulated data). *Q* is most closely related to sequences *S*_*1*_ and *S*_*2*_ along its entire genome. (TIF)

**S2 Fig.**
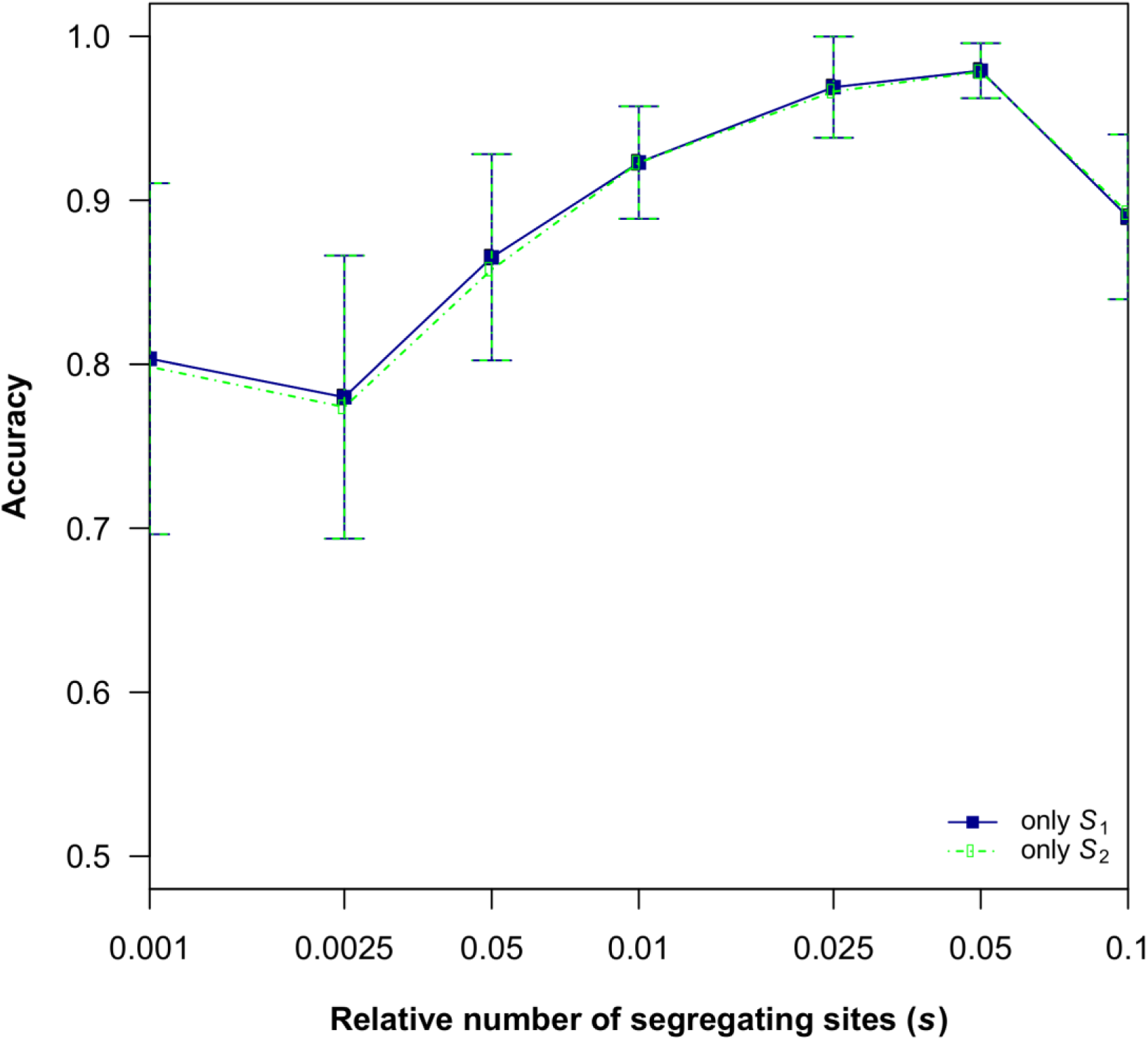
gmos accuracy when subject sequences *S*_*1*_ and *S*_*2*_ are equally closely related to a query. Proportion of correctly returned subject sequences *S*_*1*_ and *S*_*2*_ using the simulated genealogies (S1 Fig) as a function of the evolutionary distance (the relative number of segregating sites), *s*. (TIF)

**S1 File. Comparison of gmos to ClonalOrigin on simulated data sets**

The results of accuracy and speed comparison of gmos to ClonalOrigin on simulated data sets. (PDF)

**S2 File. Comparing results of gmos to the results of Palmer *et al.* [20]**.

Tables listing the genes of clade B origin along the supercontigs 1.2, 1.3, and 1.5 of *Enterococcus faecium* 1,231,408. Tables B and C also include the ClonalOrigin results from the same data set.

Table A) Genes of clade B origin along the supercontig 1.2.

Table B) Genes of clade B origin along the supercontig 1.3.

Table C) Genes of clade B origin along the supercontig 1.5. (XLS)

